# Investigation of USP30 inhibition to enhance Parkin-mediated mitophagy: tools and approaches

**DOI:** 10.1101/2021.02.02.429344

**Authors:** Eliona Tsefou, Alison S. Walker, Emily H. Clark, Amy R. Hicks, Christin Luft, Kunitoshi Takeda, Toru Watanabe, Bianca Ramazio, James M. Staddon, Thomas Briston, Robin Ketteler

## Abstract

Mitochondrial dysfunction is implicated in Parkinson disease (PD). Mutations in Parkin, an E3 ubiquitin ligase, can cause juvenile-onset Parkinsonism probably through impairment of mitophagy. Inhibition of the de-ubiquitinating enzyme USP30 may counter this effect to enhance mitophagy. Using different tools and cellular approaches, we wanted to independently confirm this claimed role for USP30. Pharmacological characterization of additional tool compounds that selectively inhibit USP30 are reported. The consequence of USP30 inhibition by these compounds, siRNA knockdown and overexpression of dominant-negative USP30 in the mitophagy pathway in different disease-relevant cellular models was explored. Knockdown and inhibition of USP30 showed increased p-Ser65-ubiquitin levels and mitophagy in neuronal cell models. Furthermore, patient-derived fibroblasts carrying pathogenic mutations in Parkin showed reduced p-Ser65-ubiquitin levels compared to wild-type cells, levels that could be restored using either USP30 inhibitor or dominant-negative USP30 expression. Our data provide additional support for USP30 inhibition as a regulator of the mitophagy pathway.

## Introduction

Mitochondrial homeostasis is important for the survival of healthy cells. Indeed, mitochondrial dysfunction has been linked to several diseases including neurodegenerative disorders such as Parkinson’s disease (PD) (Pickles et al., 2018, Calkins et al., 2011, Rodolfo et al., 2018, Clark et al., 2021). PD is a chronic, progressive neurodegenerative disease that has been linked mechanistically and genetically to alterations in mitochondrial homeostasis (Grünewald et al., 2019, Schapira et al., 1989, Bender et al., 2006, Pickles et al., 2018). It is thought that damaged mitochondria accumulate in neuronal cells, leading to neurotoxicity. In PD the dopaminergic neurones in the substantia nigra display increased neurotoxicity accompanied by mitochondrial dysfunction, leading to the characteristic motor dysfunction (Zarow et al., 2003). Thus, it has been proposed that enhanced autophagic removal of damaged mitochondria by a process called mitophagy may be a therapeutic approach for treatment of PD (Miller and Muqit, 2019).

The hypothesis that mitophagy enhancement could benefit PD was substantiated by the finding that mutations in key components of the mitophagy pathway such as *PINK1* or *PRKN* (Parkin) can cause familial autosomal recessive Parkinsonism (Kitada et al., 1998, Valente et al., 2004). Mitophagy is a multi-step process that ensures the removal of damaged mitochondria (e.g. those with loss of membrane potential) through the stabilisation of the PTEN-induced kinase 1 (PINK1) in the outer mitochondrial membrane to result in phosphorylation of PINK1 targets, most notably ubiquitin and the ubiquitin-like domain (Ubl) in Parkin at homologous serine 65 residues (Mcwilliams et al., 2018). Phospho-ubiquitin (p-Ser65-Ub) recruits and activates the Parkin E3 ubiquitin ligase, resulting in ubiquitination of target proteins such as translocase of outer mitochondrial membrane 20 (TOM20) and mitofusin-2 (Mfn-2) at the mitochondria outer membrane (Sarraf et al., 2013). Ubiquitinated mitochondria in turn recruit mitophagy receptors such as optineurin (OPTN) and nuclear dot protein 52 (NDP52) to the site of mitochondrial damage (Lazarou et al., 2015), engaging the autophagy machinery to initiate the formation of autophagosomes. Subsequently, the autophagosome engulfs damaged mitochondria and delivers them to the lysosome for degradation.

PINK1/Parkin mediated mitophagy has been proposed as a pathway containing modulatable targets for drug discovery strategies aimed at enhancing mitophagy in various diseases, including ageing and PD (Durcan and Fon, 2015, Sun et al., 2016, Stead et al., 2019). In addition to PINK1 and Parkin, de-ubiquitinating (DUB) enzymes that regulate the stability of key mitophagy regulators have also been suggested as possible drug targets (Padmanabhan et al., 2019). Several DUB’s have been shown to regulate mitochondrial homeostasis, including USP8 (ubiquitin-specific peptidase 8) (Durcan et al., 2014), USP15 (Cornelissen et al., 2014), USP30 (Bingol et al., 2014), USP33 (Niu et al., 2020) and USP35 (Wang et al., 2015).

Among these, USP30 has been investigated in depth and knockdown of USP30 was demonstrated to overcome defects in Parkin activity and enhance survival of dopaminergic neurones (Bingol et al., 2014). USP30 is localized in the mitochondrial outer membrane and was originally believed to act as the DUB that antagonises the PINK1/Parkin pathway (Bingol et al., 2014, Liang et al., 2015). However, recent findings have suggested that USP30 acts at earlier stages, possibly as a gatekeeper for mitochondrial ubiquitination, acting to dampen Parkin activity and also to prevent the unscheduled initiation of mitophagy and keep the system under tight control (Marcassa et al., 2018). It has also been suggested that a key role for USP30 is to facilitate and quality control-check protein import in mitochondria (Ordureau et al., 2020, Phu et al., 2020). Recently, it has also been shown that USP30 can also regulate peroxisome function (Marcassa et al., 2018).

The USP30 reports so far have used certain pharmacological tools, different genetic approaches and various cell lines. We wanted to conduct independent studies, searching for complementary pharmacological agents and approaches to test effects of USP30 perturbation on mitophagy in additional and different cell types. USP30 effects on mitophagy may also be dependent on the various methods to induce mitophagy. Here, several pharmacological USP30 inhibitors were characterised for their potency and selectivity against a panel of DUBs. The most selective compound was further characterised for its ability to modulate PINK1/Parkin-mediated mitophagy. Tandem Ubiquitin Binding Entities (TUBEs), p-Ser65-Ub and mitoKeima were determined in different cell models including iPSC-derived neurones and patient-derived fibroblast with pathogenic Parkin mutations. We report the results of our investigations below.

## Results

### Knockdown of USP30 enhances mitoKeima signal in SHSY5Y cells

It has been proposed that inhibition of USP30 could be a potential mechanism enabling discovery of therapeutic agents for PD (Marcassa et al., 2018, Bingol et al., 2014, Padmanabhan et al., 2019). Here, we explored USP30 inhibition via siRNAs and its effect when mitophagy was induced via different mitochondrial toxins. We used a pool of two different USP30 siRNAs (siUSP30) to achieve a greater than 80% decrease in the protein levels of USP30 in SHSY5Y cells (Figure 1A and B). The expression of USP30 protein was not affected by the addition of CCCP (Figure 1A and B). Next, we used SHSY5Y cells that stably express the mitoKeima reporter to investigate the effect of USP30 knockdown on mitophagy. After 7 days of USP30 knockdown, cells were treated with mitophagy inducers CCCP, antimycin A/oligomycin (A/O) and valinomycin (Val). In cells treated with a non-targeted (NT1) siRNA, incubation with CCCP, A/O and Val for 7 hours resulted in 2.6-fold, 1.7-fold and 2.7-fold increase in the mitoKeima signal (Figure 1C). In the USP30 knockdown cells treated with the same mitochondrial toxins a further 2-fold statistically significant increase was observed (Figure 1C). USP30 knockdown, in our hands, does not seem to significantly increase basal levels of mitoKeima signal as reported previously (Marcassa et al., 2018).

**Figure 1:**
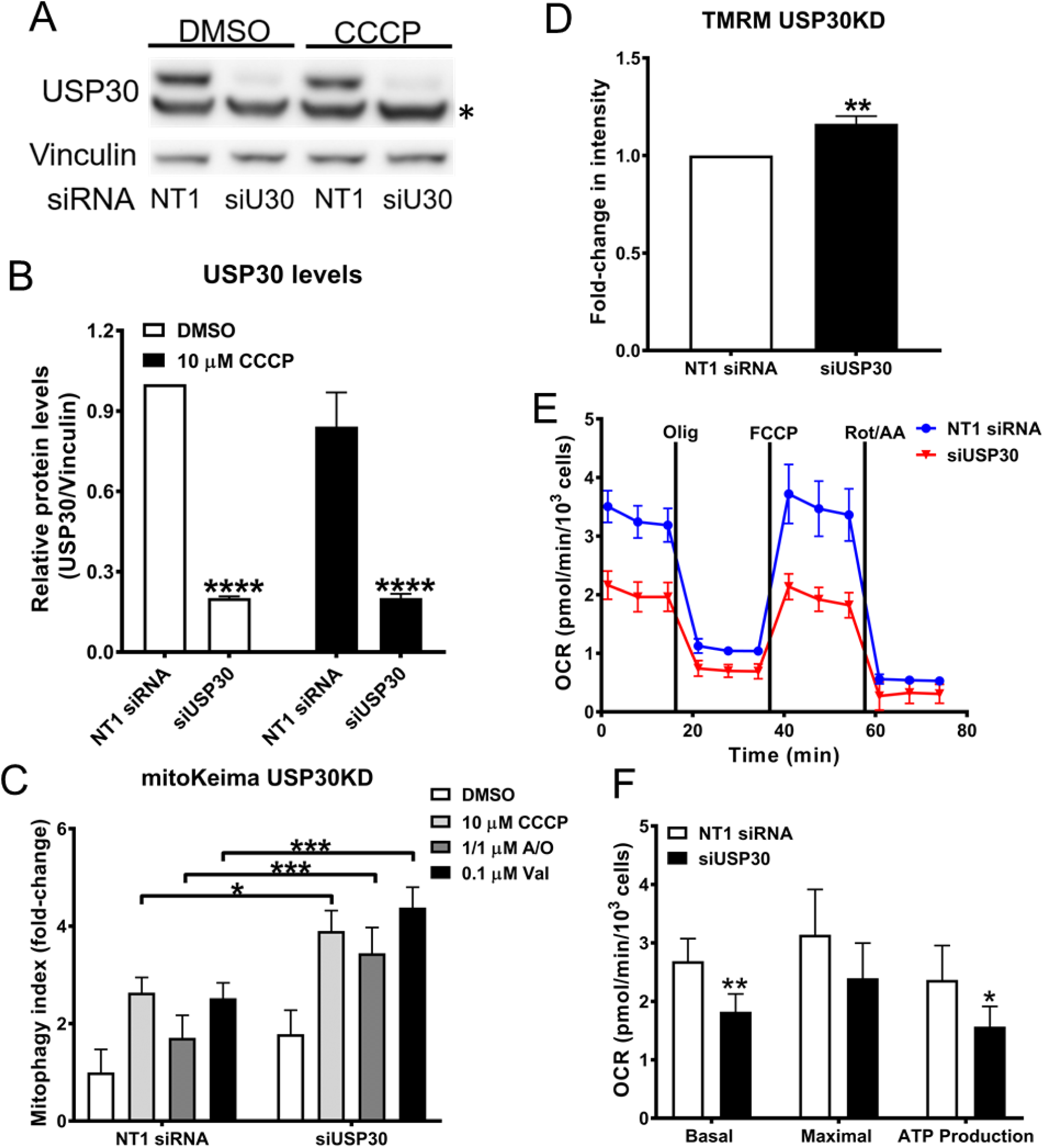
siRNA-mediated knock down of USP30 enhances mitoKeima signal in SHSY5Y cells. **(A)** Representative immunoblotting of USP30 expression levels after 7 days transfection with siUSP30 (siU30) and a non-targeted (NT1) siRNA in SHSY5Y mitoKeima cells. Vinculin was blotted for as a loading control. In parallel, incubation with 10 μM CCCP for 4 hours was tested in order to determine if it affects USP30 protein expression in the SHSY5Y mitoKeima cells. * indicate a non-specific band coming from the used USP30 antibody. **(B)** Quantification of USP30 protein expression levels from 3 independent immunoblotting experiments as presented in (A). **(C)** USP30 was knockdown in SHSY5Y mitoKeima cells for 7 days before inducing mitophagy with 10 μM CCCP or 1 μM Antimycin A/ 1 μM Oligomycin (A/O) or 0.1 μM Valinomycin (Val) for 7 hours. Cells were imaged with the Opera Phenix. With all treatments, enhanced mitophagy was observed under the USP30 knockdown conditions. The mitophagy index was calculated as the fold change of the ratio of the total lysosomal red mitochondria area divided by the total cytoplasmic green mitochondria area from 3 independent experiments. **(D)** USP30 was knockdown in SHSY5Y cells for 7 days before incubation with 50 nM TMRM and imaging with the Opera Phenix. Fold change in TMRM intensity from 3 independent experiments was quantified. **(E)** Representative OCR trace as measured by the Seahorse™ analyser in SHSY5Y cells where USP30 was knockdown for 7 days with siUSP30. **(F)** Basal respiration, maximal respiration and ATP production were determined based on the OCR measurement from 4 independent experiments. Data are pooled from 3-4 independent experiments. Error bars show means± SD. *p < 0.05, **p < 0.01, ***p < 0.001. Data were analysed with Two-way ANOVA with Sidak’s test for C and D and unpaired t-Test for D.

The effects of USP30 knockdown in SHSY5Y cells on mitochondrial function were assessed by measuring the mitochondrial inner membrane potential using TMRM and analysing mitochondrial respiration parameters. Knockdown of USP30 induced a small but statistically significant increase in TMRM (Figure 1D) and a decrease in the oxygen consumption rate (OCR; Figure 1E). Knockdown of USP30 significantly decreased basal respiration as well as oxygen consumed during oxidative phosphorylation (+ Oligomycin; Figure 1F). Reduced respiration is in agreement with a recent study in primary hepatocytes from USP30 knockout mice (Gu et al., 2020). In conclusion, we showed that genetic knockdown of USP30 can increase mitophagy as indicated by the increased mitoKeima signal but may also affect distinct aspects of mitochondrial function in the SHSY5Y cells.

### Identification of potent and selective USP30 Inhibitors

Next, we aimed to recapitulate the effects we observed with knockdown of USP30 by using a pharmacological approach. In order to accomplish that three structurally related small molecule USP30 inhibitors (which we term USP30Inh-1, -2 and -3) were synthesised based on compound structures published in patents: WO 2016/156816 and WO 2017/103614 (Figure 2A). Here, the cyano-amide functional group in the compounds is anticipated to form a covalent linkage with the catalytic cysteine within the USP30 active site, thereby inactivating the enzyme.

**Figure 2:**
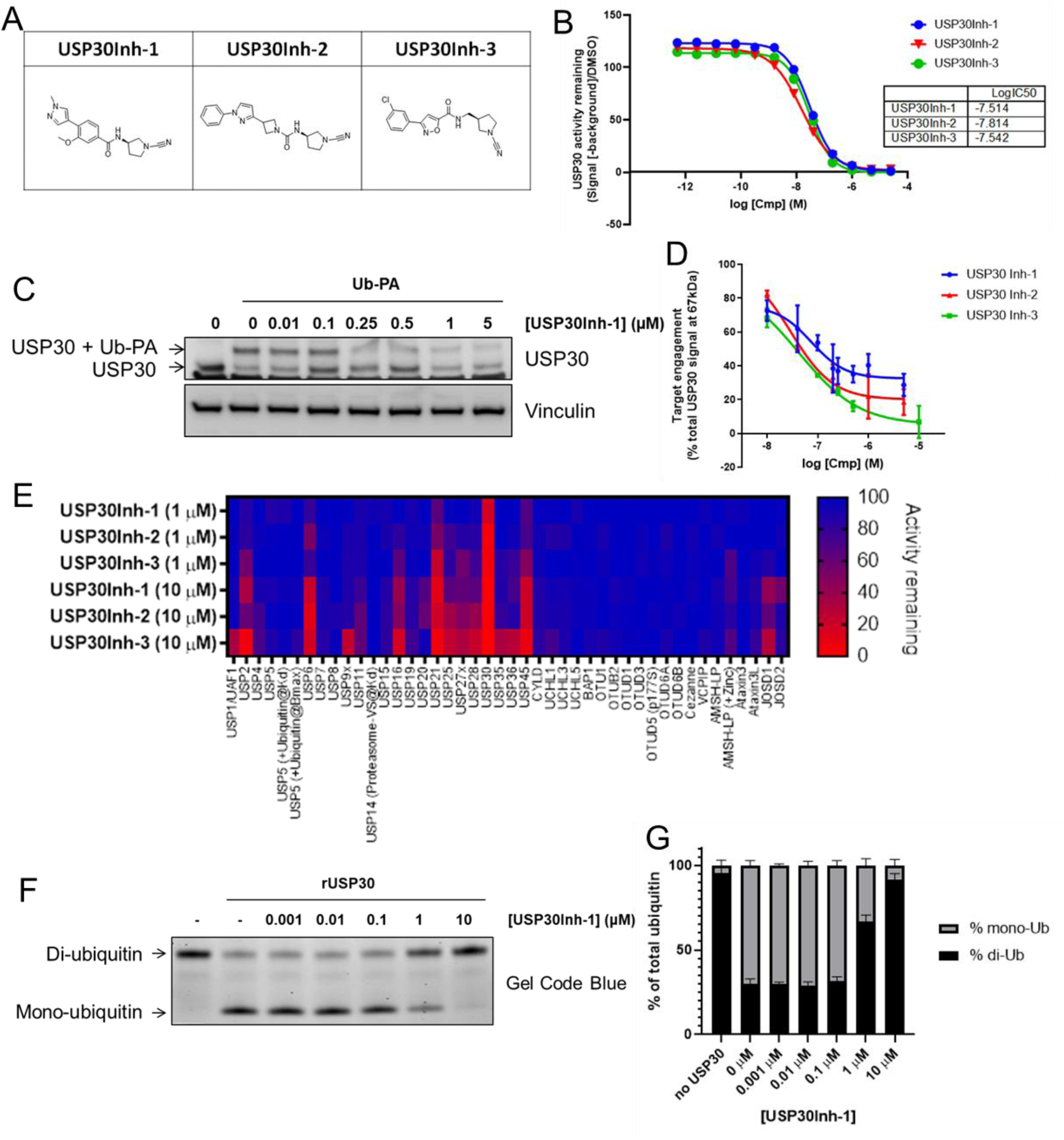
Identification of potent and selective USP30 Inhibitors. **(A)** Chemical structure of USP30Inh-1, 2 and 3. **(B)** Inhibition of the activity of recombinant human USP30 (rhUSP30) protein was tested by using 100 nM Ub-Rho110 as substrate when incubated with the indicated concentrations of USP30Inh-1, 2 and 3. **(C)** The Biotin-Ahx-Ub-PA was used with the ABP assay in order to determine in-cell target engagement of USP30 inhibitors. Representative immunoblotting for the USP30 expression when SHSY5Y cells were treated with the indicated USP30Inh-1 concentrations for 24 hours before incubating cell lysates with 2.5 μM Ub-PA for 1 hour at room temperature. Engagement of the probe is indicated by an ∼ 8 kDa shift in the molecular weight of USP30. **(D)** Quantification of the USP30 target engagement for USP30Inh-1, 2 and 3 from 2-4 independent experiments (USP30Inh-1, n=4; USP30Inh-2, n=3 and USP30Inh-6, n=2). **(E)** DUB selectivity assay (DUBprofiler™) was conducted using 1 and 10 μM of USP30Inh-1, 2 and 3. **(F)** The effect on the mono-Ub formation were assessed via SDS-PAGE where the indicated USP30Inh-1 concentrations were incubated for 2 hours at 37°C with 450 nM rhUSP30 and 2.5 µM K6-di-Ub. **(G)** Changes in in the di- and mono-Ub formation were quantified from 3 independent experiments.

**Figure 3:**
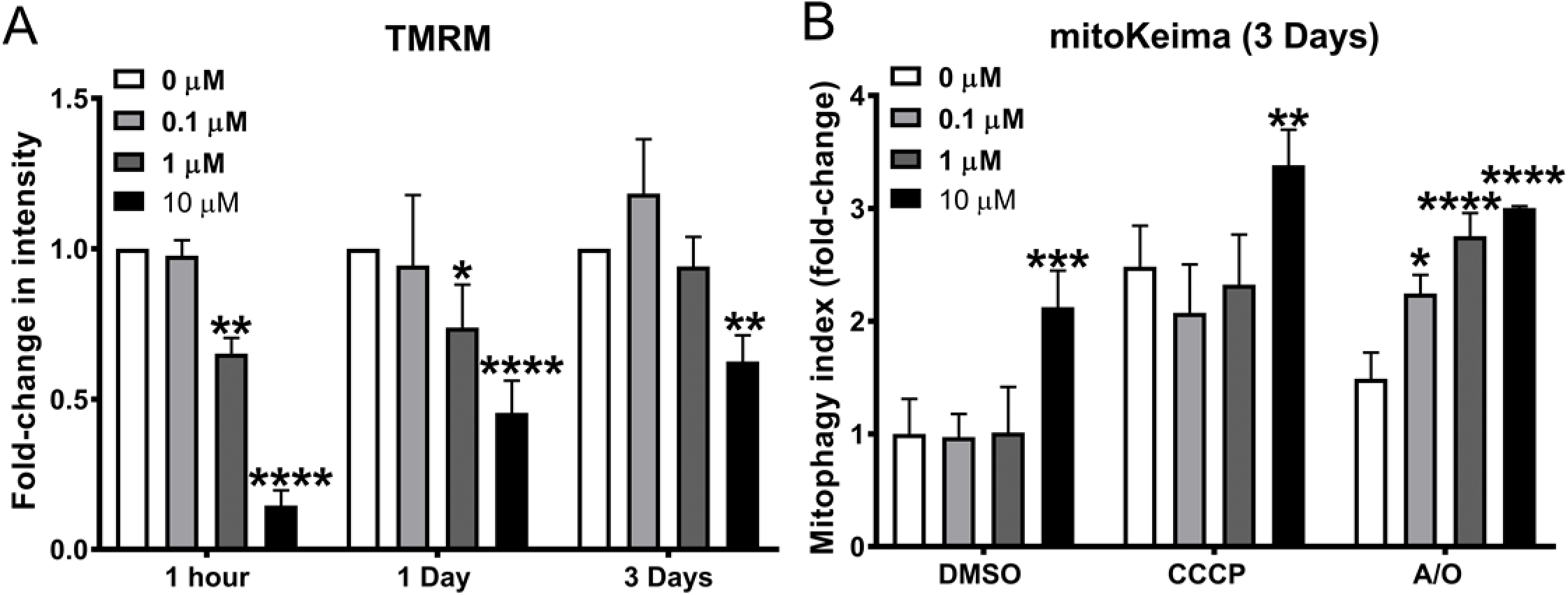
Pharmacological inhibition of USP30 enhances mitoKeima signal in SHSY5Y cells. **(A)** SHSY5Y cells were treated with the indicated USP30Inh-1 concentration for 1 hour, 1 day and 3 days. On the assay day fresh media containing the tested concentration as well as 50 nM TMRM was added and live images where acquired by the Opera Phenix. Fold change in TMRM intensity from 3 independent experiments was quantified. **(B)** SHSY5Y mitoKeima cells were treated with the indicated USP30Inh-1 concentration for 3 days. On the assay day, media was replaced and fresh medium containing fresh compound before inducing mitophagy with 1 μM A/O or 10 μM CCCP and live images were acquired with the Opera Phenix for a further 10 hours. The mitophagy index was calculated as the fold change of the ratio of the total lysosomal red mitochondria area divided by the total cytoplasmic green mitochondria area from 3 independent experiments. Data are pooled from 3 independent experiments. Error bars show means± SD. *p < 0.05, **p < 0.01, ***p<0.00, ****p<0.0001. Data were analysed with Two-way ANOVA with Sidak’s test.

USP30 inhibitory activity was assessed biochemically using the fluorogenic artificial DUB substrate, ubiquitin-rhodamine 110 (Ub-Rho110) and recombinant USP30. USP30Inh-1, -2 and -3 all potently inhibited USP30-mediated cleavage of Ub-Rho110, with calculated IC_50_ values of between 15-30 nM (Figure 2B). To determine in-cell target engagement of USP30 inhibitors, the activity-based ubiquitin probe (ABP), Biotin-Ahx-Ub-propargylamide (PA) was used. The C-terminal PA electrophile forms a covalent linkage with the active site cysteine residue of DUB enzymes. Binding of Biotin-Ahx-Ub-PA to the USP30 is observed as a band shift with an ∼8 kDa increase in USP30 molecular weight following SDS-PAGE and immunoblot analysis (Figure 2C). USP30Inh-1, -2 and -3 were observed to reduce ABP engagement in a concentration-dependent manner, suggesting compounds compete with the ABP for access to the USP30 active site catalytic cysteine, are therefore cell permeable and capable of binding endogenous USP30 (Figure 2C and D). Selectivity of USP30 inhibitors was assessed using the Ubiquigent DUBprofiler™ service. USP30Inh-1, -2 and -3 demonstrated good selectivity against >40 known DUB enzymes at 1 µM. Decrease selectivity was observed for each compound at 10 µM, with greatest off-target inhibition against USP6, USP21 and USP45 for all three compounds (Figure 2E). USP30Inh-1 demonstrated the greatest selectivity based on this panel and was taken forward for further investigation.

*In vitro*, USP30 demonstrates preference for Lys6-linked ubiquitin chains (Gersch et al., 2017, Sato et al., 2017). To confirm inhibition, we assessed USP30-mediated cleavage of Lys6-linked di-ubiquitin chains upon incubation of the recombinant USP30 with the native substrate (K6-di-Ub). In the absence of inhibitor, USP30 efficiently cleaved K6-di-Ub, to yield mono-ubiquitin (Figure 2F, lane 2). USP30Inh-1 dose-dependently inhibited USP30-catalysed cleavage of K6-di-Ub (Figure 2F and G). Together, these data confirm USP30Inh-1 as a potent and, based on the studied DUBprofiler™ panel, moderately selective, cell-permeable small molecule inhibitor of USP30. In addition to our study, Phu *et al* and Rusilowicz-Jones *et al* have also shown that compounds containing the cyano-amide functional group have some off-target activity when using higher concentrations (Rusilowicz-Jones et al., 2020, Phu et al., 2020). These studies emphasise the need to carefully profile tool compounds used in cellular studies in order to avoid effects that could be attributable to modulation of other targets.

### Pharmacological inhibition of USP30 enhances mitoKeima signal in SHSY5Y cells

In the SHSY5Y cells, the time- and concentration-dependent effect of USP30Inh-1 on mitochondrial inner membrane potential was measured using TMRM. Acute incubation (1 hour) with 10 μM USP30Inh-1 caused a depolarization of the mitochondrial membrane potential as more than 85% TMRM signal loss was observed. Increasing the incubation time to 1 and 3 days caused 40-50% depolarization at 10 μM, suggesting that the compound displays mitochondrial toxicity at this concentration. Treatment with 1 μM USP30Inh-1 also decreased TMRM after 1 hour and 1 day incubation (45% and 25% decrease, respectively). No significant effect on the mitochondrial inner membrane potential was observed after 3 days treatment with 1 μM USP30Inh-1. Incubation with 0.1 μM USP30Inh-1 had no effect in the TMRM under all tested incubation times. Therefore, we determined changes in mitophagy by measuring the mitoKeima signal in cells treated with USP30Inh-1 at the time point of 3 days. Prolonged incubation might be required in order to see changes in mitophagy after inhibition of USP30 which could be due to its suggested role in controlling the threshold of mitophagy (Rusilowicz-Jones et al., 2020, Marcassa et al., 2018).

Next, SHSY5Y mitoKeima cells were treated with 0.1, 1 and 10 μM USP30Inh-1 for 3 days. On the assay day, compound was added to the cells before inducing mitophagy by incubating cells with 1/1 μM A/O or 10 μM CCCP for 10 hours. Incubation with 10 μM USP30Inh-1 demonstrated 2-fold significant increase in basal mitoKeima signal (+ DMSO) and a further 1 to 1.4 -fold increase when CCCP or A/O was present (compared to 10 μM+DMSO). The observed changes with 10 μM USP30Inh-1 could have resulted from the mitochondrial depolarization observed using 10 μM treatment at that time point. On the other hand, cells treated with 0.1 and 1 μM USP30Inh-1, were able to significantly increase mitoKeima signal by 2.3 and 2.7-fold, respectively, but only when treated with A/O. In conclusion, we have shown that, by carefully selecting a non-toxic concentration (0.1 μM), time of incubation conditions as well as the mitochondrial toxin, we can induce mitophagy by using a potent and selective USP30 inhibitor in SHSY5Y cells.

### USP30 inhibition increases p-Ser65-Ub in dopaminergic neurones and astrocytes

To establish whether USP30Inh-1 perturbs PINK1/Parkin-mediated mitophagy in a PD-relevant cell type, the abundance of p-Ser65-Ub in iPSC-derived midbrain dopaminergic (DA) neurone/astrocyte co-cultures was assessed. p-Ser65-Ub levels are a consequence of both the activity of PINK1 and Parkin, given the specificity of PINK1-dependent ubiquitin phosphorylation and the feed-forward mechanism of Parkin recruitment and poly-ubiquitin chain elongation (Mcwilliams et al., 2018).

A large increase in p-Ser65-Ub in both tyrosine hydroxylase (TH)-positive DA neurones and glial fibrillary acidic protein (GFAP)-positive astrocytes was observed following electron transport chain (ETC) uncoupling using FCCP (Figure 4A). Following 4 days incubation with USP30Inh-1, a concentration-dependent increase in p-Ser65-Ub immunoreactivity was detected within TH-positive (Figure 4B) and GFAP-positive (Figure 4C) regions, reaching statistical significance at 1 µM. Using TMRM, a small but significant decrease in mitochondrial inner membrane potential was measured within the whole cell population but only when USP30Inh-1 reached 3 µM (Figure 4D). These data suggest that pharmacological inhibition of USP30 augments PINK1/Parkin-dependent signalling to result in increased levels of p-Ser65-Ub in this cellular system following mitochondrial uncoupling.

**Figure 4:**
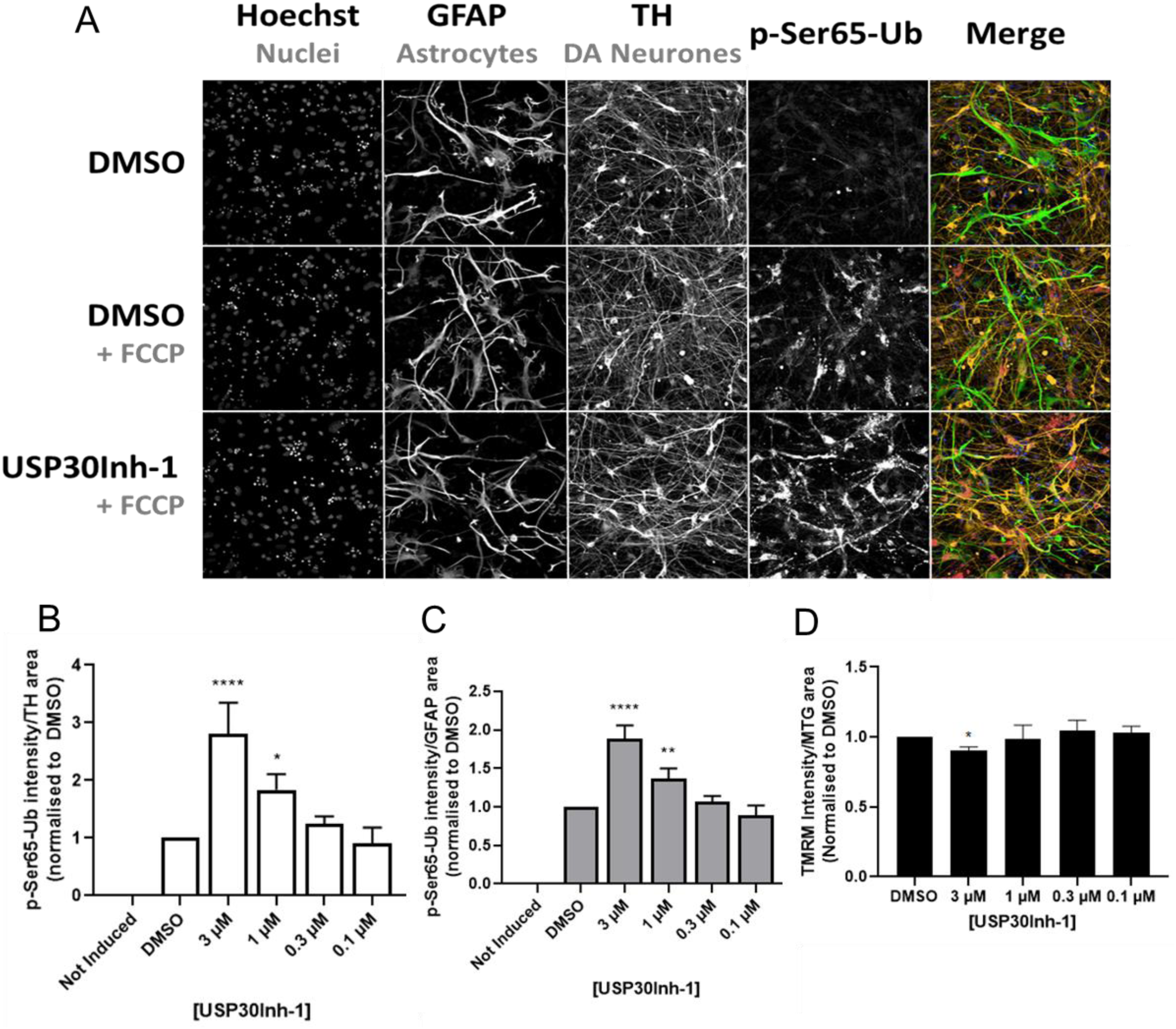
USP30 inhibition increases p-Ser65-Ub in dopaminergic neurones and astrocytes. **(A)** Representative immunostaining of p-Ser65-Ub in a co-culture of dopaminergic neurones and astrocytes. Cells were co-cultured for 5 days before incubating with 3 μM USP30Inh-1 for 4 more days. Mitophagy was induced by adding 10 μM FCCP for 2 hours. Cells were immunostained with TH, GFAP and p-Ser65-Ub. Scale bar 20 μm. **(B)** Immunoreactivity levels of p-Ser65-Ub in the dopaminergic TH-positive neurones was quantified from 3 independent experiments **(C)** Immunoreactivity levels of p-Ser65-Ub in the astrocytes-GFAP positive cells was quantified from 3 independent experiments. **(D)** The dopaminergic neurones/astrocytes co-culture were treated with the indicated USP30Inh-1 concentrations for 4 days before 50 nM TMRM/200 nM Mitotracker green (MTG) were added and imaged with the Opera Phenix. Intensity of TMRM from 3 independent experiments was quantified. Data are pooled from 3 independent experiments. Error bars show means± SD. *p < 0.05, **p < 0.01, ****p<0.0001. Data was analysed with One-Way ANOVA with Dunnett’s test.

### Fibroblasts carrying heterozygote mutations in Parkin have reduced p-Ser65-Ub levels

We next aimed to identify PD patient-derived cell lines that carry genetic defects in mitophagy signalling, whereby USP30 perturbation may have functional, disease-relevant significance. Loss-of-function mutations in *PRKN* have been shown to cause autosomal recessive, juvenile-onset PD (Kitada et al., 1998). Patient-derived fibroblasts obtained from the NINDS Biorepository were genotyped against full-length, common variant *PRKN* (NG_008289.2). Fibroblasts were selected based on zygosity at arginine 275 (R275). The Parkin R275 mutation has been linked to Parkinson’s disease through its severity of causing disruption in mitochondrial clearance (Yi et al., 2019).

Both heterozygote and compound heterozygote mutations (Parkin^+/R275W^ and Parkin^R275W/R275Q^) were confirmed using Sanger sequencing. Parkin mutations were not found within the control line (common variant; Parkin^+/+^).

Mfn2 is a well-characterised Parkin substrate (Poole et al., 2010, Gegg et al., 2010). Following induction of mitophagy, Mfn2 is rapidly extracted from the mitochondrial outer membrane and turned over by the proteasome (Chan et al., 2011). In Parkin^+/+^ fibroblasts, Mfn2 protein levels decreased, concurrent with a robust increase in Mfn2 ubiquitination after mitochondrial ETC uncoupling using FCCP. (Figure 5A, Lane 2). Increasing allelic frequency of Parkin mutation determined the degree to which Mfn2 was both ubiquitinated and degraded. Reduced Mfn2 abundance and ubiquitination was evident in the Parkin^+/R275W^ cells compared to Parkin^+/+^ (Figure 5A, Lane 5), with negligible effects observed in the Parkin^R275W/R275Q^ line (Figure 5A, Lane 8). Cellular deficits in non-selective autophagy were ruled out following appropriate processing and lipidation of LC3A/B across all genotypes in response to FCCP and chloroquine (CHL) (Figure 5A), suggesting the failure in Mfn2 processing in mutant cells was restricted to Parkin dysfunction.

**Figure 5:**
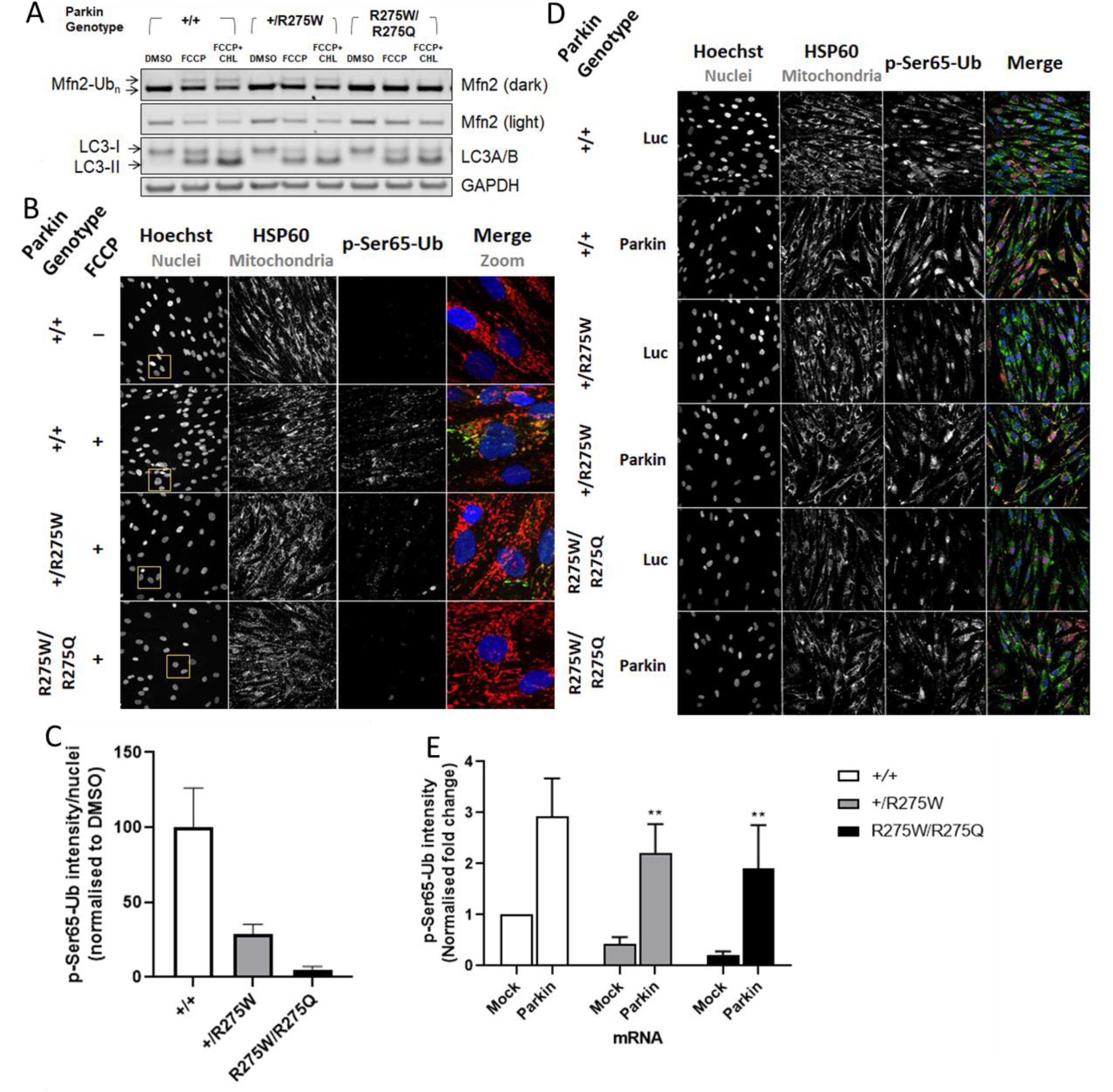
Parkin fibroblasts carrying heterozygote pathogenic mutations have reduced p-Ser65-Ub levels. **(A)** The Parkin^+/+^, Parkin^+/R275W^ and Parkin^R275W/R275Q^ fibroblasts were treated with 50 μM chloroquine (CHL) for 1 hour and then mitophagy was induced with 10 μM FCCP for further 2 hours. Expression levels of Mfn2 and LC3A/B in the different Parkin fibroblast were examined via immunoblotting **(B)** Representative images of the p-Ser65-Ub and HSP60 immunostaining in the same fibroblast as at (A). Mitophagy was induced by adding 10 μM FCCP for 2 hours. Scale bar 20 μm. **(C)** Immunoreactivity of p-Ser65-Ub was quantified from 3 independent experiment in the fibroblasts from (B) where a genetic dose-response was observed. Expression levels were normalized to DMSO Parkin^+/+^ treated cells. The Parkin^+/+^, Parkin^+/R275W^ and Parkin^R275W/R275Q^ fibroblasts were transfected with a Parkin WT mRNA and Luciferase (Luc) as a control for 20 hours before inducing mitophagy for 2 hours with 10 μM FCCP. **(D)** Representative immunostaining of the p-Ser65-Ub and HSP60 (Scale bar 20 μm) and **(E)** quantification of p-Ser65-Ub levels from 3 experiments. The expression levels were normalized to mock transfected Parkin^+/+^ cells. Data are pooled from 3 independent experiments. Error bars show means± SD. ***p < 0.001, ****p<0.0001. Data were analysed One-Way ANOVA with Dunnett’s test.

As an additional assessment of Parkin activity, p-Ser65-Ub immunoreactivity was measured and quantified (Figure 5B and C). A robust increase in p-Ser65-Ub was observed following addition of FCCP in Parkin^+/+^ fibroblasts, suggesting an active PINK1/Parkin pathway (Figure 5B). Correlating with the effects on Mfn2 (Figure 5A), a genotype-phenotype relationship, determined by allelic frequency of *PRKN* mutation was observed with respect to p-Ser65-Ub immunoreactivity in the patient-derived fibroblasts (Figure 5B and C).

We next aimed to rescue p-Ser65-Ub deficits by re-introducing functional Parkin. A full length mRNA that will express Parkin was transfected into each fibroblast line with high efficiency. p-Ser65-Ub immunoreactivity was then assessed following mitochondrial uncoupling with FCCP (Figure 5D). Re-expression of Parkin, increased p-Ser65-Ub abundance in all genotypes to similar levels (Figure 5E), suggesting the p-Ser65-Ub deficits observed are likely Parkin mediated and there are no dominant-negative effects of the mutations. Together, these data suggest Parkin mutant fibroblasts present a valuable model by which to study genetic defects in PINK1/Parkin-mediated mitophagy and to further understand disease relevance of USP30 inhibition.

### Over-expression of USP30-C77S enhances p-Ser65-Ub in Parkin heterozygote fibroblasts

We next aimed to understand the effect of USP30 perturbation on PINK1/Parkin-mediated mitophagy in Parkin compromised cells. mRNA that express the wild type or catalytically dead (dominant-negative, C77S mutation) USP30 (USP30-Myc-P2A-eGFP) was transfected with high efficiency into patient-derived Parkin mutant fibroblasts. Expression and mitochondrial localisation of both epitope-tagged WT and C77S USP30 were confirmed by co-staining with the mitochondrial marker HSP60 (Figure 6A). To confirm the C77S mutation in USP30 caused loss-of-function, USP30 activity was assessed using the ABP, Biotin-Ahx-Ub-PA. USP30-C77S failed to bind the ABP as band shift was not observed after SDS-PAGE and immunoblotting with antibody toward USP30. In contrast USP30-WT engaged the ABP, resulting in an ∼ 8kDa band shift, suggesting the WT transcript generates active enzyme. Neither WT nor catalytically dead USP30 had any effect on mitochondrial ETC subunit protein expression and robust GFP expression at ∼27 kDa was observed following all mRNA transfections, proving efficient processing of the P2A sequence (Figure 6C).

**Figure 6:**
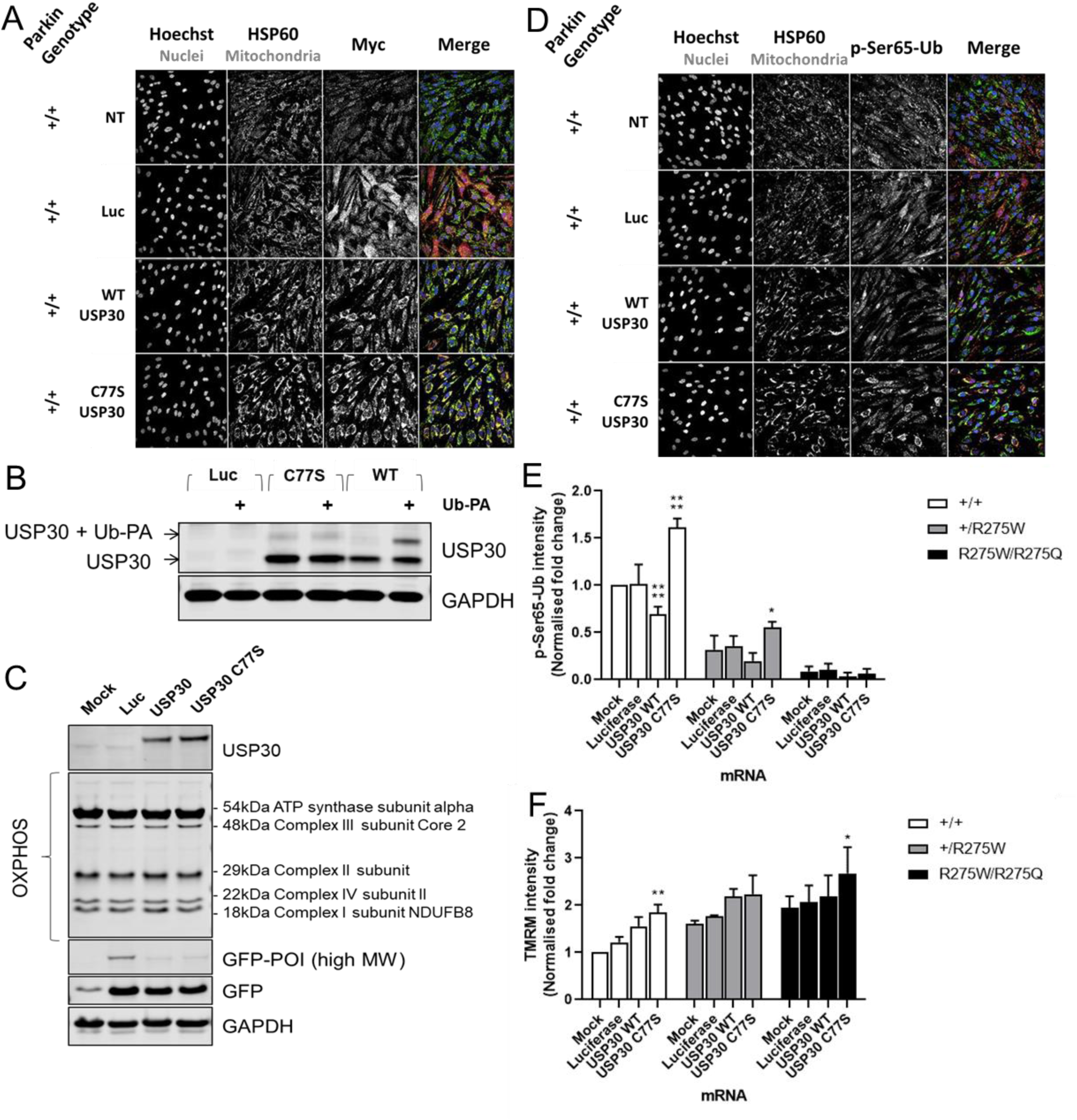
Over-expression of USP30-C77S enhances p-Ser65-Ub in Parkin heterozygote fibroblasts. **(A)** Parkin^+/+^ fibroblasts were transfected with a USP30-WT and catalytic inactive USP30 (C77S) mRNA overexpression construct for 20 hours. Expression was assessed via Immunostaining for Myc (epitope tag for USP30) and HSP60 as mitochondrial marker. Scale bar 20 μm. **(B)** Lysates from the Parkin^+/+^ fibroblasts which have been transfected with USP30-WT and USP30-C77S for 20 hours were incubated with 2.5 μM Ub-PA for 1 hour at room temperature. Engagement of probe was assessed via immunoblotting for USP30 where engagement was observed only in the cells overexpressing USP30-WT and not USP30-C77S. **(C)** Lysates from the Parkin^+/+^ fibroblasts transfected with USP30-WT or USP30-C77S for 20 hours were immunoblotted for both protein components of the mitochondrial respiratory complexes and, to determine transfection efficiency, GFP. No effect on the expression levels of the components of the mitochondrial respiratory complexes was observed under the tested conditions. The Parkin^+/+^, Parkin^+/R275W^ and Parkin^R275W/R275Q^ fibroblasts were transfected with the USP30-WT and USP30-C77S mRNA for 20 hours before conducting immunostaining for p-Ser65-Ub. **(D)** Representative images of the immunostaining for p-Ser65-Ub and HSP60 in the Parkin^+/+^ are presented. Scale bar 20 μm. **(E)** The immunoreactivity of p-Ser65-Ub was quantified 3-5 independent experiments. Expression levels were normalized to mock transfected Parkin^+/+^ cells. **(F)** Same conditions as (D) were tested but cells were incubated with 50 nM TMRM/200 nM MTG before imaging on the Opera Phenix. Intensity of TMRM from 3-5 independent experiments was quantified. Expression levels were normalized to mock transfected Parkin^+/+^ cells. Data are pooled from 3-5 independent experiments. Error bars show means± SD. *p < 0.05, **p < 0.01, ****p<0.0001. Data were analysed One-Way ANOVA with Dunnett’s test.

We next assessed the effect of USP30 overexpression on p-Ser65-Ub immunoreactivity following mitochondrial uncoupling. Expression of USP30-WT significantly decreased p-Ser65-Ub abundance compared to firefly luciferase control in Parkin^+/+^ cells (Figure 6D and E). The same trend was observed in the Parkin^+/R275W^ line; however, statistical significance of effects was not reached (Figure 6D). In contrast, expression of the USP30-C77S produced a significant increase in p-Ser65-Ub abundance in both the Parkin^+/+^ and Parkin^+/R275W^ lines (Figure 6D). No change in p-Ser65-Ub abundance was observed in the Parkin^R275W/R275Q^ (Figure 6D). TMRM analysis revealed a small apparent hyperpolarisation following USP30-C77S over-expression in Parkin^+/+^ and Parkin^R275W/R275Q^ lines (Figure 6E), potentially an imaging artefact due to observed morphological changes associated with the over-expression. Given inner membrane depolarisation is necessary to recruit and activate Parkin, we anticipate this will have little effect on p-Ser65-Ub abundance. Taken together, USP30, wild type or C77S mutant, can directionally impact p-Ser65-Ub levels, an event dependent on functional Parkin.

### Enhanced p-Ser65-Ub levels upon pharmacological inhibition of USP30 in Parkin heterozygote fibroblasts

We next determined the effects of pharmacological inhibition of USP30 in the Parkin mutant fibroblasts. Western blot analysis revealed increased p-Ser65-Ub following FCCP in both the Parkin^+/+^ and Parkin^+/R275W^ fibroblasts, and slightly enhanced following 6 days incubation with USP30Inh-1 (Figure 7A). Parkin^R275W/R275Q^ cells displayed minimal increases in p-Ser65-Ub following FCCP, with no effect of USP30Inh-1 (Figure 7A). The Parkin genotype determined the degree of Mfn2 ubiquitination, with strong mono-ubiquitination in the Parkin^+/+^ fibroblasts and absence of any ubiquitinated species in the Parkin^R275W/R275Q^ line (Figure 7A). In all genotypes, under the conditions of the experiments, no effect of USP30Inh-1 was observed on Mfn2 mono-ubiquitination.

**Figure 7:**
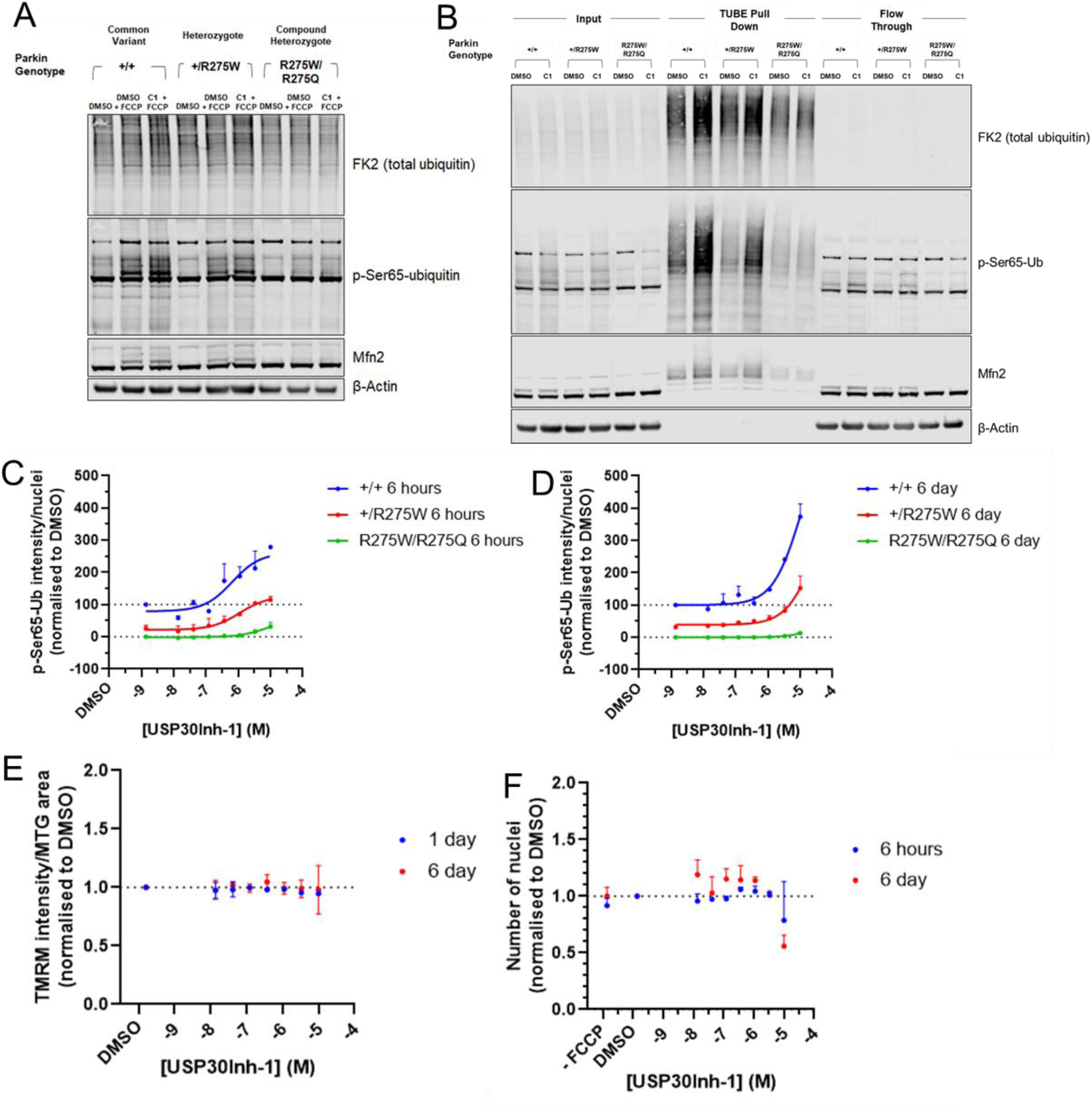
Enhanced p-Ser65-Ub levels upon pharmacological inhibition of USP30 in Parkin heterozygote fibroblasts. **(A)** The Parkin^+/+^, Parkin^+/R275W^ and Parkin^R275W/R275Q^ fibroblasts were treated with 3 μM USP30Inh-1 for 6 days before inducing mitophagy with 10 μM FCCP for 2 hours. Lysates were immunoblotted for total ubiquitin, p-Ser65-Ub and Mfn2. **(B)** Same conditions as at (A) but TUBE enrichment was used to enhance the levels of poly-ubiquitinated proteins before conducting immunoblotting. The Parkin^+/+^, Parkin^+/R275W^ and Parkin^R275W/R275Q^ fibroblasts were treated with the indicated concentrations of USP30Inh-1 for **(C)** 6 hours or **(D)** 6 days before inducing mitophagy with 10 μM FCCP for 2 hours. The immunoreactivity of p-Ser65-Ub was quantified from 3 independent experiments for each condition. The expression levels were normalized to DMSO Parkin^+/+^ treated cells (+/+= 100% and Parkin^R275W/R275Q^ = 0%). **(E)** The Parkin^+/+^ fibroblasts were treated with the indicated concentrations of USP30Inh-1 for 1 day or 6 days. On the assay day, cells were incubated with 50 nM TMRM/200 nM MTG before imaging on the Opera Phenix. Intensity of TMRM from 3 independent experiments was quantified and no change was observed. **(F)** The Parkin^+/+^ fibroblasts were treated with the indicated concentrations of USP30Inh-1 for 6 hours or 6 days. Cells were counted after staining for nuclei and average from 3 independent experiment was calculated. Reduced numbers of nuclei and therefore cells was observed only at 10 μM USP30Inh-1 after 6 days incubation.

To further understand ubiquitination dynamics in response to USP30 inhibition, ubiquitin pull-down was performed using TUBEs. TUBEs enrich for poly-ubiquitinated proteins, with lower affinity for mono-ubiquitinated (Lopitz-Otsoa et al., 2012) and a clear increase in total poly-ubiquitin can be detected following TUBE pull down (Figure 7B). In the presence of FCCP, poly-ubiquitinated species of Mfn2 were observed with TUBE pull-down in both Parkin^+/+^ and Parkin^+/R275W^ lines, the greatest levels found in the Parkin^+/+^ cells (Figure 7B). In both Parkin^+/+^ and Parkin^+/R275W^ fibroblasts, the Parkin genotype-dependent increase in Mfn2 poly-ubiquitination was further enhanced with 6 days treatment of USP30Inh-1 (Figure 7B). In the Parkin^R275W/R275Q^ fibroblasts, Mfn2 poly-ubiquitination was not enhanced following incubation with USP30Inh-1 (Figure 7B). We identified FCCP/USP30Inh-1-dependent increases in Mfn2 and p-Ser65-Ub correlated with Parkin genotypes.

To confirm our observations and understand the kinetics associated with the ubiquitination response, Parkin mutant fibroblasts were incubated with USP30Inh-1 for either 6 hours or 6 days and levels of p-Ser65-Ub were assessed following FCCP treatment. As observed in Figure 5C, a Parkin genotype-dependent response was observed in p-Ser65-Ub immunoreactivity. At both 6 hours and 6 days incubation, a USP30Inh-1 concentration-dependent increase in p-Ser65-Ub immunoreactivity was observed, correlating with severity of Parkin mutation (Figure 7C and D). In the Parkin^+/+^ fibroblasts, USP30Inh-1-dependent changes were greater following 6 days incubation and independent of any changes in mitochondrial inner membrane potential as measured using TMRM (Figure 7E). A small increase in p-Ser65-Ub was observed in the Parkin^R275W/R275Q^ fibroblasts, potentially suggesting some remaining functionality in Parkin. Notably, with 6 days treatment at 10 μM USP30Inh-1 but not at lower concentrations, a ∼40% decrease in cell number, was observed in the Parkin^+/+^ fibroblasts (Figure 7F), potentially reflecting increased cell death, cell detachment or decreased cell proliferation. Taken together, USP30 inhibition can, under appropriate conditions with careful selection of concentrations and time of incubation, rescue ubiquitination deficits in patient-derived Parkin^+/R275W^ fibroblasts to the levels of Parkin^+/+^. Functional Parkin is therefore required for USP30 inhibition to positively modulate the mitophagy cascade.

## Discussion

Mitochondrial dysfunction is a prominent pathological feature of both sporadic and familial PD (Bose and Beal, 2016, Hasson et al., 2013, Grünewald et al., 2019). PINK1 and Parkin are sensor and amplifier proteins integral to removing damaged mitochondrial by mitophagy (Park et al., 2006, Clark et al., 2006, Narendra et al., 2008). The association between mutations in PINK1 and Parkin and the development of PD, suggests that defective mitophagy and accumulation of damaged mitochondria are key factors involved in the aetiology of disease. In addition to genetic deficits implicated in mitophagy, mitochondrial dysfunction and reduced rates of mitophagy are evident in sporadic PD (Hsieh et al., 2016, Ambrosi et al., 2014, Smith et al., 2016), again linking mitochondrial health and clearance processes to PD pathophysiology. Oxidative stress and bioenergetics failure are recognised phenotypes of PD *in vivo* and *in vitro* (Grünewald et al., 2010, Langley et al., 2017) and improving the efficiency of mitochondrial quality control will likely prevent inappropriate generation of mitochondrially-derived oxidative intermediates and cellular damage (Miller and Muqit, 2019). Therefore, enhancement of mitochondrial clearance by mitophagy has been proposed as a disease-modifying strategy in PD. USP30 has been suggested as a potential target for enhancing mitophagy since it is the main DUB that is localised in the outer mitochondrial membrane. Studies have demonstrated knockout of USP30 to enhance mitophagy in both cellular and animal models. (Padmanabhan et al., 2019, Bingol et al., 2014, Liang et al., 2015, Marcassa et al., 2018). In PINK1 or Parkin knockout *Drosophila* models, USP30 knockout was able to protect from a PD-like motor phenotype (Bingol et al., 2014).

Initially, we confirmed that siRNA-mediated knockdown of USP30 increases stress-induced mitophagy. We utilised the neuroblastoma SHSY5Y cell line and showed that knockdown of USP30 could be demonstrated to enhance the mitoKeima signal indicating changes in the mitophagy pathway, in agreement with the other studies (Bingol et al., 2014). Knockdown of USP30, however, seems to affect the function of the mitochondrial ETC since a decrease in basal respiration and oxygen used for ATP synthesis was determined by the OCR measurement which was also accompanied by an increase in the mitochondrial membrane potential (hyperpolarization). To generate ATP, mitochondria utilise the proton electrochemical gradient potential generated by serial reductions of electrons in the ETC. The reductive transfer of electrons through the ETC complexes provides the force to drive protons from the mitochondrial matrix to outer face of the inner mitochondrial membrane against their concentration gradient. The accumulated protons in the inter membrane space then flow back into the mitochondrial matrix through Complex V (ATP synthase), producing ATP. These reactions result in the formation of a voltage and a pH gradient (Mitchell, 1966). Studies have shown that mitochondrial hyperpolarization can be caused by inhibition of Complex I and Complex V of the ETC (Forkink et al., 2014, Solaini et al., 2007). Inhibition of these complexes could then lead to reduced OCR levels. The effect of USP30 knockdown in the OCR and inner mitochondrial membrane, assuming no off-target effects could possibly be attributed to the newly characterised role of USP30 in regulating protein import in the mitochondria (Phu et al., 2020, Ordureau et al., 2020). Phu *et al* has recently shown that a number of proteins belonging to the ETC (such as ATPB, NDUA6 and COX4-1) presented increased ubiquitination when USP30 was inhibited (Phu et al., 2020). The absence of these proteins from the inner mitochondrial membrane could result in a compromised ETC and that could lead to reduced respiration. Another study by Gu *et al* has reported reduced OCR in primary hepatocytes from USP30 knockout mice where they identified that USP30 can be phosphorylated by IKKβ, resulting in deubiquitination of ATP citrate lyase and fatty acid synthase (Gu et al., 2020). Both our results and the study from Gu *et al* indicate that USP30 could potentially regulate different metabolic pathways such as ATP production and lipogenesis.

The observed results with the genetic perturbation of USP30 were replicated with the small molecule inhibitor USP30Inh-1 resulting in a dose dependent increase in the mitoKeima signal when mitophagy was induced with A/O. However, in the SHSY5Y cells, 10 μM USP30Inh-1 caused a depolarization of the mitochondrial inner membrane potential when cells were treated for 1 hour. Prolonged incubation seems to restore the mitochondrial membrane potential whereas lower concentrations (0.1 μM) do not seem to affect it. A small but significant decrease in TMRM was also observed in the DA neurones/astrocytes co-cultures when treated with 3 μM USP30Inh-1 for 4 days. The acute drop in the TMRM signal after 1 hour incubation with 10 μM USP30Inh-1 could mean that the cells were metabolically affected by pharmacological inhibition of USP30 and/or other targets of USP30Inh-1. Based on the DUB profiling treatment with 10 μM USP30Inh-1, where we see the highest toxicity, seems to target other DUBs (e.g. USP21 and USP45), which indicates that the compound could potentially have other intracellular targets. A significant decrease in mitochondrial potential could indicate that at higher concentration USP30Inh-1 is acutely toxic to cells by possibly affecting the enzymes of the ETC that are responsible for the maintenance of the inner membrane potential, in line with a suggested role for USP30 in mitochondrial protein import (Phu et al., 2020, Ordureau et al., 2020). For transient effects on TMRM, a compensation mechanism may take place to result in partial restoration of the mitochondrial inner membrane potential. Nonetheless, we are able to show that a non-toxic concentration (0.1 μM) could also increase the mitoKeima signal.

The USP30 inhibitors used here are proposed to form an adduct between the cyano-amide group and the cysteine residue in the active side of the protein. Recently, two other USP30 inhibitors were reported containing the cyano-amide group. Phu *et al* reported USP30i which was able to increase TOM20 ubiquitination at concentrations above 5 μM in HEK293 cells overexpressing Parkin (Phu et al., 2020). The second study comes from Rusilowicz-Jones *et al*, where they characterised the compound FT385 which increased TOM20 ubiquitination, mitolysosomal formation via monitoring the mitoQC signal and p-Ser65-Ub levels at 200 nM in SHSY5Y cells (Rusilowicz-Jones et al., 2020). There has only been limited assessment of overall mitochondrial function for these compounds, and our study suggests that careful evaluation of effects on mitochondrial health is important when considering applications in cell biology. Equally, compounds such as USP30Inh-1 have not been completely characterised and it is possible that they have effects and targets beyond those explored in this study.

Further, we evaluated genetic and pharmacological inhibition of USP30 in a PD-relevant model. Following mitochondrial damage, both Parkin and ubiquitin are phosphorylated by PINK1 on Ser65 residues (Kondapalli et al., 2012, Kane et al., 2014). These phosphorylation events result in full activation of Parkin E3-ligase activity and translocation to mitochondria, tagging them for removal by the autophagosome-lysosome system (Wauer et al., 2015, Kazlauskaite et al., 2015, Narendra et al., 2008). The R275W mutation within the RING1 domain of Parkin is one of the most common autosomal recessive, early onset-associated variants (Bayne and Trempe, 2019). The R275W mutation has limited effect on Parkin recruitment to depolarized mitochondria but causes severe disruption of subsequent mitochondrial clearance (Geisler et al., 2010, Yi et al., 2019). Interestingly, a low level of mono-ubiquitinated VDAC1 has been detected with Parkin R275W, suggesting remaining, residual enzymatic functionality (Geisler et al., 2010). These observations are consistent with small depolarisation-induced increases in Mfn2-Ub and p-Ser65-Ub in the Parkin^R275W/R275Q^ fibroblasts (Figure 7).

Parkin comprises of an N-terminal Ubl domain and four RING-like domains (RING0, RING1, IBR and RING2). Upon activation, conformational change results in rearrangements of these domains to allow coordination of the p-Ser65-Ub molecule. The Ubl domain interacts with helix 261–274 within the RING1 region, served by hydrogen bonds between Asp274 immediately preceding the Arg275 mutation site (Sauvé et al., 2015). The Arg275 mutation site interacts with Glu321 in the p-Ser65-Ub -binding helix, and the R275W mutation has been proposed to cause clashes within the p-Ser65-Ub -binding helix and with regions of the Ubl domain (Yi et al., 2019). Together, these structural insights predict the Arg275 mutation to lead to Parkin destabilisation, and are consistent with observations of reduced Parkin protein expression and cellular aggregation (Yi et al., 2019, Hampe et al., 2006). Here, we have characterised patient-derived familial PD fibroblasts with verified R275 mutation in Parkin. In agreement with structural and functional insights, heterozygous and compound homozygous mutant fibroblasts demonstrate: mitophagy deficits (but normal autophagic flux); reduced Mfn-2 ubiquitination and clearance; and decreased p-Ser65-Ub, with a clear genotype-phenotype relationship determined by allelic number of *PRKN* R275 mutations. Furthermore, re-introduction of common variant Parkin protein can rescue mono- and bi-allelic mutation at R275, restoring p-Ser65-Ub levels to that of control fibroblasts.

Considering both the proposed role of USP30 in controlling initiation of mitophagy and the structural and functional elements of the Parkin R275W mutation, we focused upstream within the PINK1/Parkin cascade to explore and identify biochemical changes associated with genetic and small molecule USP30 inhibition in this context. To date, PINK1 is the only recognized ubiquitin kinase (Kane et al., 2014, Schubert et al., 2017); consequently, Ser65 phosphorylation of ubiquitin is a key biomarker of cellular efficacy in recognition and clearance of dysfunctional mitochondria. Following mitochondrial uncoupling, small molecule USP30 inhibition was able, under appropriate conditions, to rescue Parkin-dependent deficits in mitophagy signalling, restoring p-Ser65-Ub and ubiquitinated Mfn2 toward Parkin^+/+^ levels. Notably, this phenotype was clearly observed in Parkin^+/R275W^ fibroblasts but not in the Parkin^R275W/R275Q^. Equally, expression of a catalytically dead version of USP30 was able to phenocopy pharmacological USP30 inhibition with respect to increasing mitochondrial p-Ser65-Ub. These data strongly suggest a requirement for functional Parkin in mediating USP30-dependent ubiquitination dynamics.

In this report, we have explored pharmacological and genetic inhibition of USP30 in multiple cell models. We have: (1) demonstrated that the PINK1-Parkin-USP30 axis is conserved across cell types (DA neurones, astrocytes, SHSY5Y and fibroblasts); (2) expanded the USP30 inhibitor toolbox by characterizing an additional small molecule inhibitor of USP30 that increases the mitoKeima signal in SHSY5Y cells when using non-toxic concentrations, consistent with the effects of genetic experimental perturbation of USP30; (3) characterised a panel of patient-derived fibroblasts carrying Parkin mutations, demonstrating defects in Parkin function; (4) recognised that genetic or pharmacological USP30 inhibition can rescue Parkin-dependent signalling defects and (5), shown that Parkin activity is necessary for USP30 inhibitor-driven events during the early steps of mitophagy initiation.

Our findings indicate that USP30 may have a significant role in regulating Parkin-mediated mitophagy as well as ubiquitination and the PINK1-dependent phosphorylation of ubiquitin. Genetic inhibition of USP30 was able to rescue Parkin-dependent deficits in the mitophagy pathway. However USP30 seems also to have an important role in the ETC function since its genetic inhibition caused a metabolic changes. The pharmacologic inhibition of USP30 presented off-target effects as well as increased toxicity at the highest tested concentrations indicating that more specific and possibly chemical diverse compounds are required. Further studies will be required to explore the role of USP30 in mitochondrial function and metabolic homeostasis.

### Limitations of the Study

In the current study we characterised new small molecule compounds that can dose-dependently inhibit USP30. However, these compounds seem to present some off-target effects since inhibition of at least two other DUBs was observed as well as a time-dependent effect on the mitochondrial membrane potential. We also observed that genetic inhibition of USP30 seems to affect the mitochondrial function. Further studies will be required in order to explore the effect of both pharmacological and genetic perturbation of USP30 in mitochondrial metabolism.

### Resource Availability

Further information as well as data that support these finding are available upon request to the Lead Contact.

## Methods

### Materials and Compound synthesis

All chemicals and compounds were purchased from Sigma-Aldrich unless otherwise specified. USP30 inhibitors were derived from patent ID: WO 2016/156816 and WO 2017/103614 and synthesised in-house.

### Antibodies

The following antibodies were used for western blotting: anti-Mfn2 (Abcam, ab56889, 1 in 500, anti-total ubiquitin (FK2, Enzo, BML-PW8810-0100, 1 in 500), anti-phospho-Ser65-ubiquitin (Millipore, ABS1513-I, 1 in 300), anti-GAPDH (CST, 2118L, 1 in 2000), anti-OXPHOS cocktail (Abcam, ab110411, 1 in 500), anti-LC3A/B (CST, 12741T, 1 in 1000), anti-VDAC1 (Abcam, ab154856, 1 in 1000), anti-USP30 (Sigma, HPA016952, 1 in 500), anti-USP30 (Santa Cruz, sc-515235, 1 in 100), anti-Vinculin (Abcam, ab129002, 1 in 10000), anti-GFP (ab13970, 1 in 1000))

The following antibodies were used for immunocytochemistry: anti-phospho-Ser65-ubiquitin (Millipore, ABS1513-I, 1 in 1000), anti-phospho-Ser65-ubiquitin (CST, 62802, 1 in 1000), anti-HSP60 (Abcam, ab128567, 1 in 1000), anti-tyrosine hydroxylase (Millipore, AB1542, 1 in 100), anti-Beta III tubulin (Biolegend, 801201, 1 in 2000), anti-GFAP (CST, 3670S, 1 in 200).

### Cell culture

All cell culture media and supplements were purchased from Fisher Scientific unless otherwise specific. SHSY5Y (human neuroblastoma) cells were purchased from ATCC and cultured in Dulbecco’s Modified Eagle Medium (DMEM; high glucose plus GlutaMAX), supplemented with 10% fetal bovine serum (FBS) and 100 U/ml penicillin/100 µg/ml streptomycin (Gibco). Parkin mutant fibroblasts were obtained from the NINDS Repository Parkin^+/+^ control (ND36320), Parkin^+/R275W^ (ND29369) and Parkin^R275W/R275Q^ (ND40072) were grown in DMEM (high glucose, plus GlutaMAX), containing 10% fetal calf serum (FCS; Labtech), 1 mM sodium pyruvate and 100 U/ml penicillin/100 µg/ml streptomycin.

iDOPA neurones and iAstrocytes (both Cellular Dynamics International) were plated on a matrix of poly-l-ornithine (Sigma, 0.01%) and laminin (Sigma, 10 µg/ml in D-PBS) in clear bottom 96 well Cell Carrier Ultra plates (PerkinElmer) at a density of 100,000 and 10,000 cells per well, respectively. Cells were left to mature over 5 days, feeding every 2-3 days before compound treatment.

### USP30 activity assay

USP30 inhibitors were dispensed into black, clear bottom, low binding 384 well plates (Greiner) using the ECHO 550 (Labcyte) liquid handler. 75 nl of compound was dispensed in 100% DMSO. Total reaction volume was 30 µl producing an USP30 inhibitor top concentration of 25 µM. His-tagged, 2x concentrated recombinant human USP30 protein (rhUSP30; amino acids 57-517 of the full-length protein, and a C-terminal 6-His tag, Sf 21 (baculovirus)-derived human USP30 protein) (10 nM; final assay concentration = 5 nM; Boston Biochem) was prepared in USP30 activity assay buffer (50 mM Tris base, pH 7.5, 100 mM NaCl, 0.1 mg/ml BSA (Sigma, A7030), 0.05% Tween 20, 1mM DTT) and 15 µl dispensed into compound containing assay plate using the Multidrop dispenser (Thermo Scientific) and incubated for 30 min at room temperature. Following incubation, 15 µl of 2x concentrated ubiquitin-rhodamine 110 (Ub-Rho110) (200 nM, final assay concentration = 100 nM; Boston Biochem), prepared in USP30 activity assay buffer, was dispensed into the compound-rhUSP30 containing plate and fluorescence immediately read on the FLIPR TETRA plate reader (Molecular Devices). Fluorescence was recorded every 30 seconds over 1 hour and intensity analysed.

### Di-ubiquitin cleavage assay

His-tagged rhUSP30 was diluted to 600 nM in USP30 activity assay buffer (see above) and incubated with USP30Inh-1 where appropriate for 15 min at room temperature. Lysine-6-di-ubiquitin (K6-di-Ub) was solubilised to 30 µM following manufacturer’s instructions and further diluted to 10 µM in USP30 activity assay buffer. His-tagged rhUSP30-USP30Inh-1 was combined with a K6-di-Ub at a ratio 3:1, yielding final assay concentration of 450 nM and 2.5 µM respectively. The plate was centrifuged, covered and incubated for 2 hours at 37°C. The reaction was stopped following the addition of 4x sample buffer (containing 10% beta mercaptoethanol (BME; Thermo Fisher)) to each well and heated to 95°C for 10 min. Proteins were resolved by SDS-PAGE, gel was fixed (50% methanol, 7% acetic acid in water) for 15 min and wash 3 times in water. GelCode Blue (Fisher Scientific) was used to stain proteins for 1.5 hours to overnight, de-stained using water and imaged using the Li-Cor Odyssey CLx. Mono- and di-ubiquitin band intensity was quantified using Image Studio (Licor).

### DUB selectivity profiling

Selectivity profiling of 41 DUB enzymes was performed at Ubiquigent (Dundee, U.K.) using the DUBprofiler™ platform and Ub-Rho110-glycine substrate based-assay.

### SDS-PAGE and western blot

Cells were washed in cold D-PBS prior to lysis on ice using lysis buffer (1% NP40, 100mM Tris pH8, 100mM NaCl, 10% glycerol, 5mM EDTA, cOmplete™ EDTA-free Protease Inhibitor Mixture, Roche Applied Science) supplemented with 20 mM N-ethylmaleimide. Lysates were cleared by centrifugation at 15,000 x g at 4 °C, and the resulting pellet was discarded. Total protein concentrations were determined using Pierce BCA protein assay kit (Thermo Fisher Scientific), and lysates were diluted to approximately equal concentrations before addition of 4x sample buffer (containing 10% BME) with immediate boiling at 95°C for 10 min. Proteins were separated by 4-12% bis-tris SDS-PAGE (Invitrogen) in 1x MOPS or MES SDS running buffer (Invitrogen) at 100V, and transferred (1x Transfer Buffer (Invitrogen), 20% methanol) onto Immobilon-FL PVDF membrane (Millipore) for 1.5 hours at 100V. Membranes were blocked in Odyssey blocking buffer TBS (Licor) or 5% non-fat milk or 1% BSA (Sigma, A7906) for 1 hour before over-night incubation at 4°C with primary antibodies. Membranes were scanned the following day after 1 hour incubation with secondary antibodies using Odyssey CLx Imaging System or ImageQuant system (BioRad, HRP anti-rabbit or anti-mouse (CST, 7074S/ 7076S) were used as secondary antibodies).

### Target engagement using activity-based probe (ABP); Biotin-Ahx-Ub-PA

SHSY5Y cells were plated at 0.5× 10^6^ cells per well in 6 well plates. Following compound incubation, cells were washed 3 times in ice-cold D-PBS (without Ca^2+^ and Mg^2+^) and lysed in HR buffer (50 mM Tris base, 250 mM sucrose, 5 mM MgCl_2_, 0.1% NP40, 0.5% CHAPS) with DTT (1 mM) and 1x HALT™ Protease and Phosphatase Inhibitor Cocktail (Fisher Scientific) added immediately before lysis. Cell lysates were sonicated 3 times using a probe tip sonicator (5 seconds; amplitude = 6, Soniprep 150 Plus, MSE) and centrifuged at 4°C for 10 min at 18,000x *g*. Supernatants were collected and protein quantified by BCA assay and diluted to 1 mg/ml in HR lysis buffer.

Biotin-Ahx-Ub-PA (UbiqBio) was solubilised to 100 µM following manufacturer’s instructions. An equivalent of 2.5 µM ABP per 100 µg protein lysate was incubated at room temperature for 1 hour with constant agitation. Assay was terminated following addition of 4x concentrated sample buffer (Licor), containing 10% BME. Samples were heated to 95°C for 10 min before separation by SDS-PAGE and western blotting.

### Ubiquitinated protein enrichment using TUBEs

Fibroblasts were plated at 1× 10^6^ cells per 10 cm dish and left to adhere over-night. The following day, USP30Inh-1 was added where appropriate and replenished with a full media change at day 3. At day 6, mitophagy was induced using FCCP (10 µM) for 2 hours. Plates were washed twice in ice-cold D-PBS (without Ca^2+^ and Mg^2+^) and lysed in 300 µl NP40 lysis buffer (1% NP40, 100 mM Tris base pH 8, 100 mM NaCl, 10% glycerol, 5 mM EDTA, final concertation pH 8). 1x HALT™ Protease and Phosphatase Inhibitor Cocktail, 50 µM PR-619 and 10 mM N-ethylmaleimide (Fisher Scientific) were added immediately before lysis. Cells were scraped and homogenates collected.

Lysates were centrifuged at 4°C for 10 min at 18,000x *g* and supernatant collected and input sample removed. 50 µl of Magnetic TUBE1 (LifeSensors) was added to 300 µl of cell lysate and suspension left to rotate at 25 rpm over-night at 4°C. The following day, samples were placed on a magnetic rack, flow through sample removed. Remaining lysate was removed, and magnetic beads washed 4 times with ice-cold Tris buffered saline-0.1% Tween 20 (TBS-T). Following the final wash, protein were eluted using 40 µl of 1x Licor sample buffer (containing 2.5% BME) and heated for 10 min at 95°C. Proteins were resolved by SDS-PAGE and western blotting.

### Immunocytochemistry (ICC)

Cells washed with D-PBS and ice-cold fixing solution (1:1; acetone: methanol) was added, and plates incubated for 2 min on ice. Cells were washed twice using D-PBS and permeabilised using 0.25% Triton X (Bio-Rad) in D-PBS for 20 min. Cells were blocked for 1 hour at room temperature (3% BSA (Sigma, A3803) in D-PBS). Primary antibodies diluted in blocking buffer, added to cells and plates were incubated at 4°C over-night. Cells were washed twice in TBS-T and secondary antibody solution was added: anti-mouse Alexa-488 and anti-rabbit Alexa-647 (Fisher Scientific) diluted 1/1000; nuclear stain (Hoechst 33342, final concentration of 2 µg/ml) diluted in blocking buffer and incubated for 1 hour at room temperature. Cells were washed twice in TBS-T and imaged using PerkinElmer Opera Phenix.

### Measurement of mitophagy levels by using the mitoKeima reporter

For the Keima reporter assay, SHSY5Y cells were transfected with the mitochondria-targeted monomeric Keima-Red (mitoKeima) (Medical and Biological laboratories Co., Ltd. AM-V0251) (Soutar et al., 2019, Katayama et al., 2011). Stable cell lines were generated by culturing cells in media containing 500 μg/ml G418 (Sigma). For treatments with USP30Inh-1, cells were seeded into 384 well Cell Carrier Ultra plates (PerkinElmer). The following day cells were treated with different concentrations of USP30Inh-1 for 3 days. At the assay day, media was replenished with DMEM lacking phenol red (Thermo Scientific, containing 10% FBS, 100 U/ml penicillin/100 µg/ml streptomycin and 1x L-Glutamine) which contained fresh compound and then mitopahgy was induced by adding (final concentrations) 10 μM CCCP or 1/1 μM A/O. Two μg/ml Hoechst 33342 (Thermo Fisher) was included in order to identify nuclei. Images were acquired on the PerkinElmer Opera Phenix high content confocal microscope using the 63x water objective with temperature and CO_2_ controls enabled. Excitation wavelengths and emission filters used were as follow: Cytoplasmic Green Keima: 488 nm, 650–760 nm; Lysosomal red Keima: 561 nm, 570–630 nm; Hoechst: 375 nm, 435–480 nm. Images were analysed by using a CellProfiler pipeline as follows: Cells, cytoplasmic green and lysosomal red mitoKeima areas were segmented and the area occupied by the two respective mitoKeima signals was determined. The mitophagy index was calculated as the ratio of the total lysosomal red mitochondria area divided by the total cytoplasmic green mitochondria area.

### Transfection

For mRNA transfection in fibroblasts mRNA’s were synthesised *de novo* (Trilink Biotechnologies) using cDNA sequences for *USP30* (accession NM_032663) and introducing point mutation c.G230C p.C77S and *PRKN* (accession NM_004562.3). 7 µg mRNA was transfected into 0.5× 10^6^ fibroblasts using Cell Line Nucleofector Kit V (Lonza) and AMAXA program X-001 following the manufacturer’s instructions. Cells were assayed 18-20 hours later and EGFP expression was confirmed by imaging.

For USP30 knockdown in SHSY5Y cells was conducted as previously described (Marcassa et al., 2018) by following the “double hit” approach where cells transfected twice over a period of 7 days. Briefly, cells were seeded in either 12-well plates (0.1× 10^6^ cells) or 96-well CellCarrier Ultra plates (8,000 cells, PerkinElmer) before transfecting with 40 nM ONTARGETplus Non-targeting oligo siRNA (NT1 siRNA: 5’-UGGUUUACAUGTCGACUAA-3’, Dharmacon) or with a pool of two different USP30 siRNAs (D1: 5’-CAAAUUACCTGCCGCACAA-3’; D3, 5’-ACAGGAUGCUCACGAAUUA-3’, Dharmacon; siUSP30) by using Lipofectamine RNAiMAX (Thermo Scientific) following the manufacturer’s instructions.

### Oxygen consumption rate (OCR) measurement

Mitochondrial function in SHSY5Y cells was measured by using the Seahorse XFe96 Extracellular Flux Analyser (Agilent). For treatment with USP30Inh-1 8,000 cells per well were seeded in the Seahorse 96 well plate and left to attach overnight. Next day, cells were treated with different USP30Inh-1 concentrations for 24 hours. On the assay day, cell culture media was washed and replaced with fresh assay media (XF basic media; Agilent), supplemented with 10 mM Glucose, 2 mM L-Glutamine and 1 mM Sodium Pyruvate (pH was adjusted to 7.4). The plate was incubated at 37°C for equilibration for 1 hour before loading to the analyser. Mitochondrial respiration was measured by using the Mito-Stress Test (Agilent) as per manufacturer’s instructions. Oligomycin (1 μM), FCCP (1.2 μM), rotenone (1 μM) and antimycin A (1 μM) were sequentially added to cells to determine mitochondrial respiration parameters. For the normalization step, 1 μg/ml Hoechst 33342 (Thermo Fisher) was added to the cells and incubated for 10 min before imaging on the PerkinElmer Opera Phenix. Regarding the measurement of OCR in the USP30 knockdown in SHSY5Y cells, cells were initially transfected in 6-well plates by following the forward transfection approach (see above) for the “first hit” whereas for the “second hit” a reverse transfection approach was followed. Briefly, for the reverse transfection, reactions containing diluted Lipofectamine RNAiMAx and 40 nM of the respective siRNAs were incubated for 20 min at room temperature. In the meantime, cells transfected for 3 days were trypsinised, counted and diluted in order to obtain 15,000 cells in 41.5 μl OptiMEM per well before seeding in the Seahorse 94 well plate. Consequently, 8.5 μl of the siRNA reaction was added per well. Cells were incubated for 7 hours before replacing with fresh media. At the 7 day transfection time point, mitochondrial respiration was measured as described above.

### Mitochondrial membrane potential

Mitochondrial membrane potential was measured using TMRM in re-distribution mode. Media was replaced with compound containing TMRM staining solution (50 nM tetramethyl rhodamine methyl ester (TMRM), 200 nM Mitotracker Green FM (MTG; Fisher Scientific), 2 µg/ml Hoechst) in complete cell culture media and left for 1 hour at 37°C to equilibrate. Images were acquired using the Perkin Elmer Opera Phenix with temperature and CO_2_ controls enabled.

### Data analysis and statistics

Data are presented as mean±standard deviation (SD). Normalisation of the data allowed for control of inter-assay variability. Curve fitting was performed using GraphPad Prism version 6.05 for Windows. Statistical tests are indicated in figure legends. Statistical significance was assessed as being p<0.05.

## Acknowledgements

This work was primarily funded by the UCL:Eisai Therapeutic Innovation Group under the TIGE project. Additional funding for CL and RK was from the Michael J Fox Foundation, Medical Research Council funding to the MRC LMCB (MC_U12266B) and an MRC Confidence-in-Concept award (MC_PC_13078). The PerkinElmer Opera Phenix at the UCL High-content Biology Laboratory MRC LMCB was co-funded by the MRC Dementia Platform UK (MR/M02492X/1) and the MRC LMCB (MC_U12266B). We would like to thank members of the UCL:Eisai Therapeutic Innovation Group (Andy Takle, Tom Warner, Peter Atkinson, Hélène Plun-Favreau, Adrian Isaacs, Anthony Groom, Jane Kinghorn, Lorna Ravenhill and Ged Corbett) for scientific discussions, guidance and critique of the project. Also, for help with management and oversight of the UCL:Eisai collaboration.

## Author Contributions

Conceptualization of the study: J.S, T.B and R.K; Design and execution of experiments, data analysis, and preparation of figures; ET, A.W, E.C, A.H, C.L, B.R and T.B; Provision of chemistry support: K.T and T.W; Manuscript writing: E.T, J.S, T.B and R.K.

